# Meta-analyses identify differentially expressed microRNAs in Parkinson’s disease

**DOI:** 10.1101/253849

**Authors:** Jessica Schulz, Petros Takousis, Inken Wohlers, Ivie O G Itua, Valerija Dobricic, Gerta Rücker, Harald Binder, Lefkos Middleton, John P A Ioannidis, Robert Perneczky, Lars Bertram, Christina M Lill

## Abstract

**Objective:** MicroRNA-mediated (dys)regulation of gene expression has been implicated in Parkinson’s disease (PD), although results of microRNA expression studies remain inconclusive. We aimed to identify microRNAs that show consistent differential expression across all published expression studies in PD.

**Methods:** We performed a systematic literature search on microRNA expression studies in PD and extracted data from eligible publications. After stratification for brain, blood, and cerebrospinal fluid (CSF)-derived specimen we performed meta-analyses across microRNAs assessed in three or more independent datasets. Meta-analyses were performed using effect-size and *p*-value based methods, as applicable.

**Results:** After screening 599 publications we identified 47 datasets eligible for meta-analysis. On these, we performed 160 meta-analyses on microRNAs quantified in brain (n=125), blood (n=31), or CSF samples (n=4). Twenty-one meta-analyses were performed using effect sizes. We identified 13 significantly (Bonferroni-adjusted α=3.13×10^-4^) differentially expressed microRNAs in brain (n=3) and blood (n=10) with consistent effect directions across studies. The most compelling findings were with hsa-miR-132-3p (*p*=6.37×10^-5^), hsa-miR-497-5p (*p*=1.35×10^-4^), and hsa-miR-133b (*p*=1.90×10^-4^) in brain, and with hsa-miR-221-3p (*p*=4.49×10^-35^), hsa-miR-214-3p (*p*=2.00×10^-34^), and hsa-miR-29c-3p (*p*=3.00×10^-12^) in blood. No significant signals were found in CSF. Analyses of GWAS data for target genes of brain microRNAs showed significant association (α=9.40×10^-5^) of genetic variants in nine loci.

**Interpretation:** We identified several microRNAs that showed highly significant differential expression in PD. Future studies may assess the possible role of the identified brain miRNAs in pathogenesis and disease progression as well as the potential of the top blood microRNAs as biomarkers for diagnosis, progression or prediction of PD.

## INTRODUCTION

Parkinson’s disease (PD) is the second most common neurodegenerative disease affecting 1% of people over the age of 60. The increasing incidence of PD in industrialized, aging populations constitutes a growing socio-economic burden (1). Idiopathic PD results from a combination of multiple genetic (2-4) and environmental/lifestyle factors (5,6). However, the currently known risk factors only explain a small fraction of the phenotypic variance of PD. Likewise, PD progression and its response to therapy represent multifactorial processes that are only poorly understood (6).

It is likely that epigenetic mechanisms contribute to PD development and progression (6,7). Epigenetics refers to regulatory mechanisms of gene expression that are not mediated by the DNA sequence itself but by chemical or allosteric DNA modifications or by the action of regulatory non-coding RNAs. MicroRNAs (miRNAs) are small non-coding RNAs that serve as posttranscriptional regulators of gene expression. They bind to messenger RNA (mRNA) and promote their degradation and/or decrease their translation (8). In brain, miRNAs appear to play a role in essentially all processes related to neuronal function, including the development of neurodegenerative disorders such as PD (9-11). The prominent role that miRNAs may play for the integrity of the central nervous system is exemplified by experiments inducing a selective depletion of Dicer, the enzyme that cleaves precursor forms of miRNAs (pre-miRNAs) into mature miRNAs. Depletion of this protein in midbrain dopaminergic neurons in mice leads to neurodegeneration and locomotor symptoms mimicking PD (12). However, identifying specific miRNAs playing important roles in PD development and progression remains a challenge. In humans, several studies have reported on differential miRNA expression in PD patients compared to controls, but results have been inconclusive. This is in part due to the fact that sample sizes tend to be comparatively small and that studies often analyze different tissues or biological fluids (Table 1). As a consequence, it has become exceedingly difficult to interpret the often discrepant results.

**Table 1.**
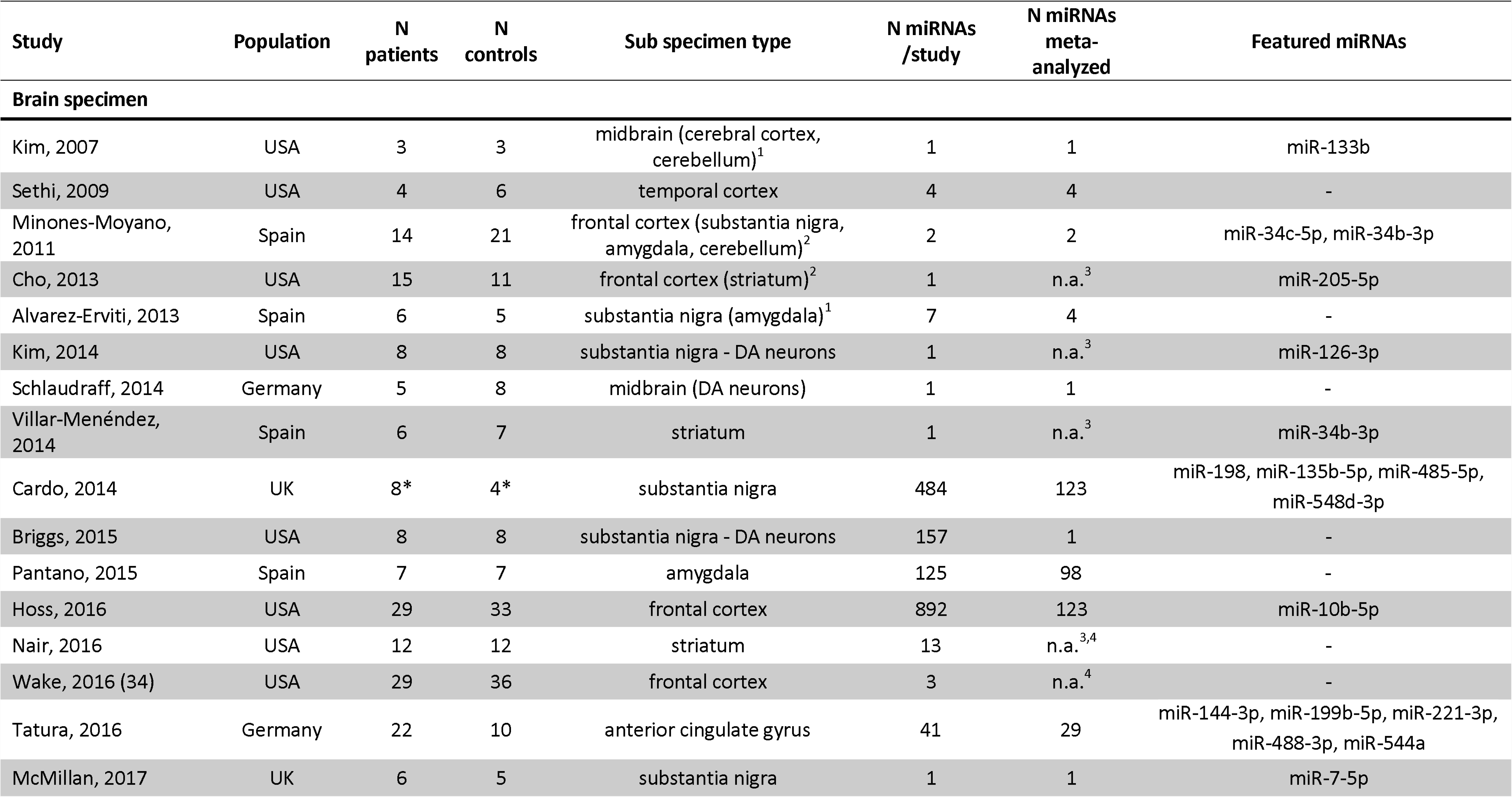
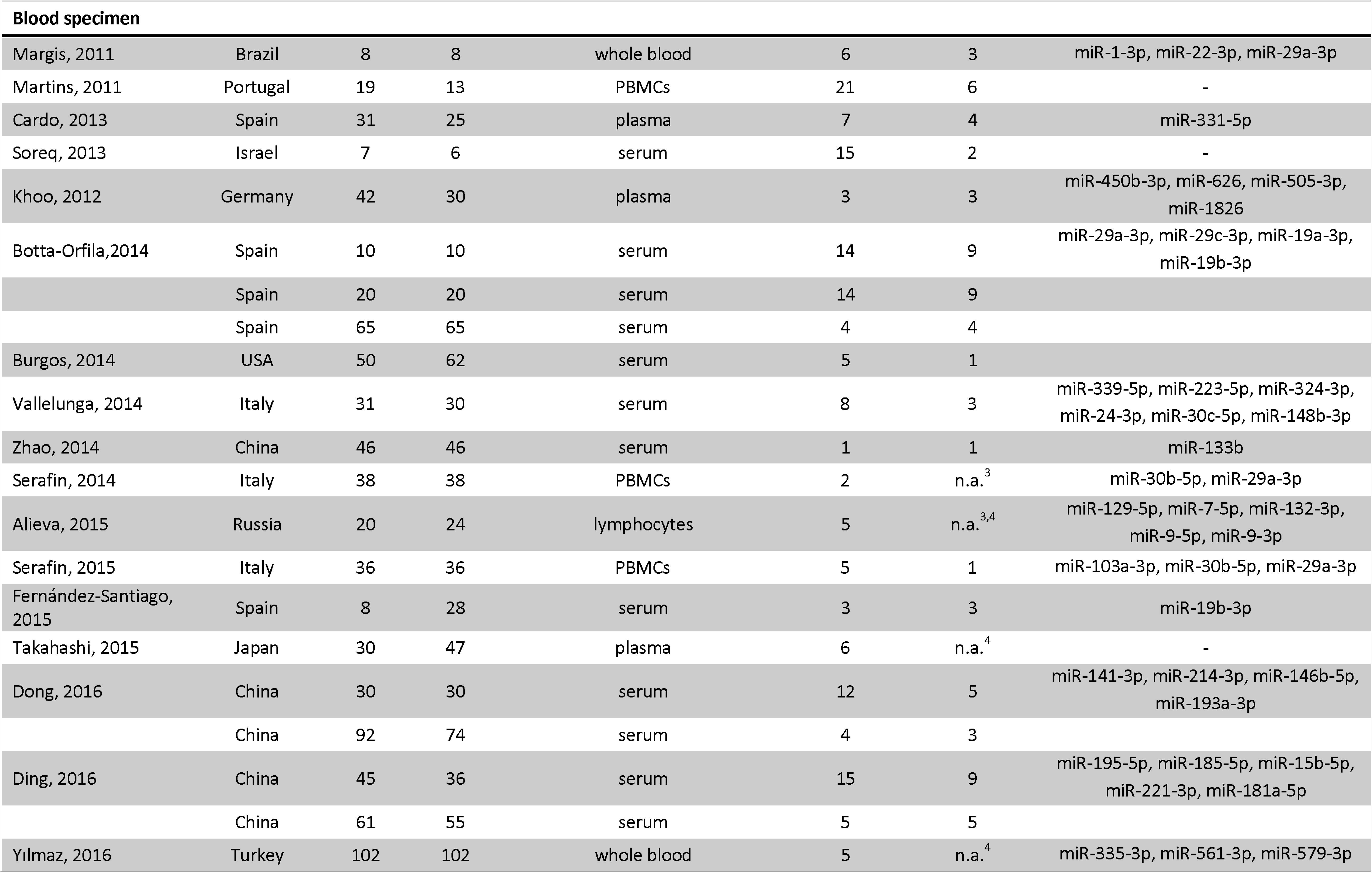
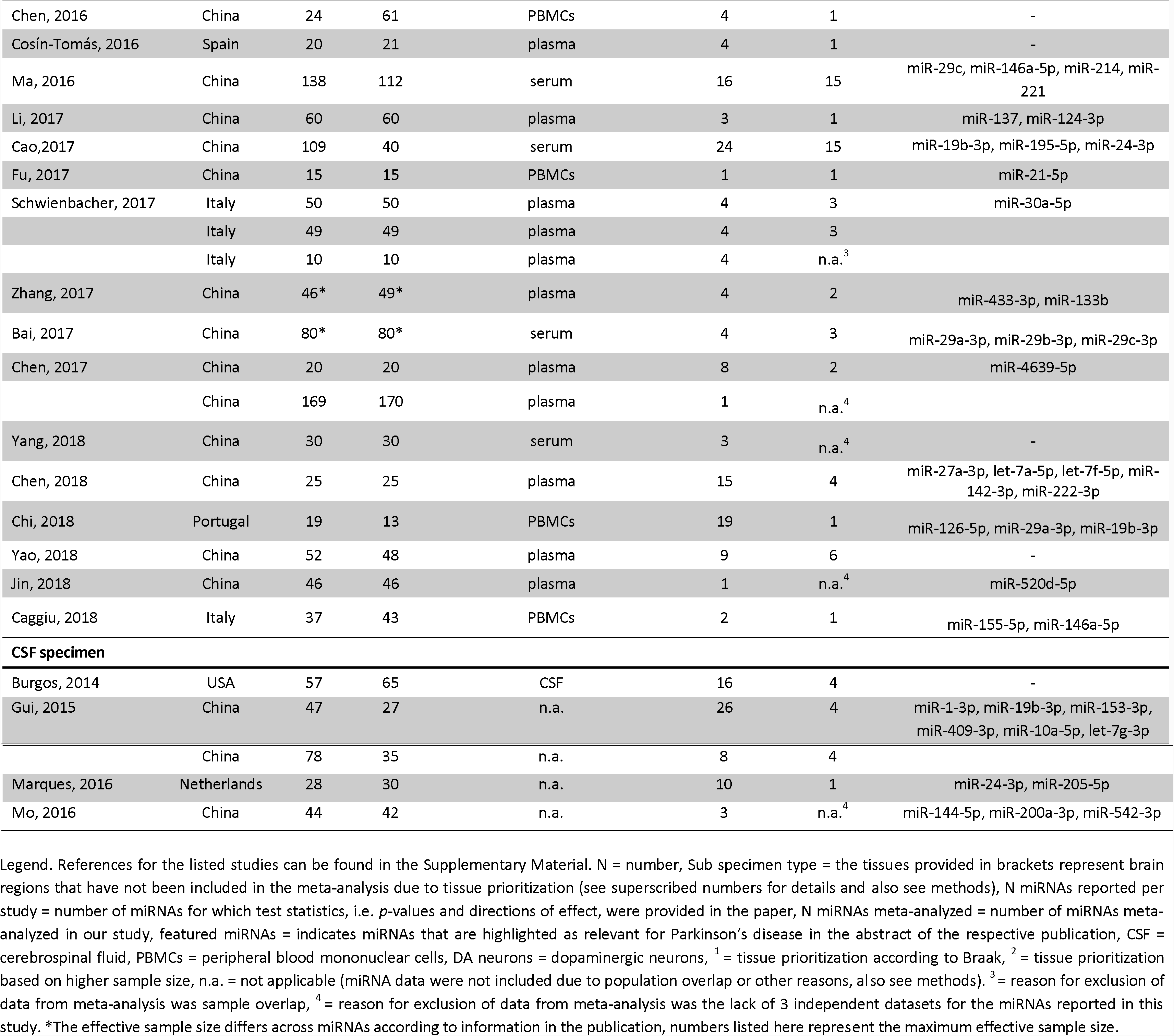
Overview of published microRNA expression studies in Parkinson’s disease patients and controls

One way to address this challenge is to assess the cumulative evidence for differential miRNA expression, e.g. by systematic meta-analyses combining all available published expression data in the field. Such approaches demonstrated their value in the context of genetic associations and environmental risk factors in several multifactorial diseases including PD (e.g. ref. (3,5)). For gene expression studies, combining published data by meta-analysis is a particularly challenging task due to the non-standardized fashion that data are reported across publications. The aim of this study was to overcome these difficulties and to identify consistently differentially expressed miRNAs in PD based on published evidence. To this end, we performed a systematic literature search to identify all relevant miRNA expression studies comparing idiopathic PD versus control subjects and extracted data from all eligible papers using a standardized protocol optimized for the extraction of expression data. Finally, we applied *p*-value based meta-analyses in order to identify miRNAs that are consistently differentially expressed in PD.

## METHODS

### Literature search and eligibility criteria

The work-flow and data collection procedures applied in this study (Figure 1) are similar to those for genetic association studies developed earlier by our group (3,13), adapted to the characteristics of gene expression studies. A systematic literature search for miRNA expression studies in PD was performed using PubMed (http://www.pubmed.gov) applying the search term “(microRNA OR miRNA OR miR* OR micro-RNA) AND Parkinson*”. Citations were assessed for eligibility using the title, abstract, or full text, as necessary. Only articles in English and published in peer-reviewed journals (last PubMed search date: October 1^st^, 2018) were considered. Original studies comparing the expression of miRNAs in patients with clinical and/or neuropathological diagnosis of PD and unaffected controls were included. Studies were included irrespective of patient treatment status. MiRNA expression studies on monogenic PD or PD families were excluded, as well as studies examining only patients with PD with dementia. A summary of eligible studies can be found in Table 1.

**Figure 1.**
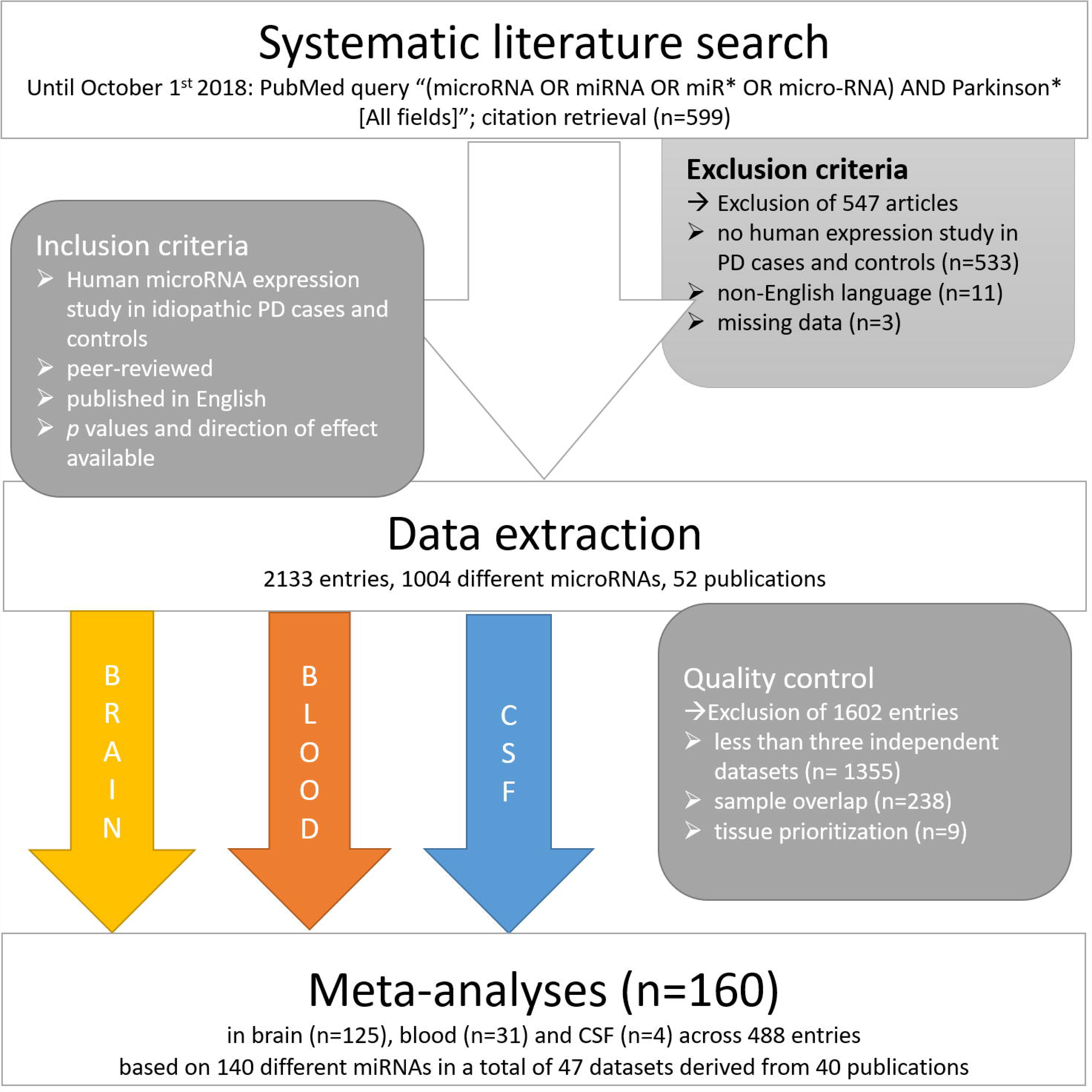
Flowchart of literature search, data extraction, and analysis of miRNA expression data. Note to the editor and reviewers: The numbers displayed in this figure have been updated along with our update of the datafreeze of our literature search, changes in the figure have not been tracked

### Data extraction

Details extracted for each eligible study consisted of the first author name, year of publication, and the PubMed identifier, along with key study- and population-specific details such as population and city of origin, number of idiopathic PD patients, number of controls, source of specimen (i.e., brain, blood, and/or CSF, and a more specific description for each specimen type, e.g. substantia nigra, frontal cortex, amygdala, etc., or whole blood, serum, PBMCs, etc.), experimental method(s) used, identifiers of the miRNAs, their expression in samples of PD patients versus controls (i.e. up- or downregulation or no difference), and corresponding *p*-values. Where available, effect-size estimates (means and standard deviations [used as provided or calculated from 95% confidence intervals or standard errors), mean differences and corresponding measures of variance] and/or details on the applied test statistics were extracted. Some of the effect-size data were extracted from results displayed in figures by using a specialized data capture program (“Plot digitizer”, http://plotdigitizer.sourceforge.net). All extracted data were double-checked by an independent member of our group against the original publications.

For quality control, we assessed reported miRNAs for their inclusion in miRBase, v21 (http://www.mirbase.org). MiRNA names corresponding to expired entries, non-human miRNAs, or non-miRNA sequences not listed in miRBase were excluded from the analysis. MiRNAs reported in the included studies were aligned to mature miRNA sequences according to miRBase. The same mature miRNA sequence reported with different miRNA names in different publications (applicable to 10/2133 entries) were subsumed under one common identifier. This concerned miRNAs hsa-miR-199a-3p/hsa-miR-199b-3p, hsa-miR-365a-3p/hsa-miR-365b-3p, hsa-miR-517a-3p/hsa-miR-517b-3p.

### Data cleaning and reformatting

Data were analyzed after stratification for specimens derived from “brain”, “blood”, and “CSF”. Potential sample overlaps, i.e. investigations of the same miRNA in identical or overlapping datasets of the same specimen type (i.e. brain, blood or CSF), for instance in two different publications, were systematically assessed in each stratum. Overlap was determined based on the origin and descriptions of the datasets, overlapping coauthors and/or references to previous studies. In case of sample overlap, only the data entry from the largest dataset was retained for further analysis. In some datasets (n=3), miRNAs were assessed in more than one brain tissue in the same (or largely overlapping) individuals. Here we chose only one brain tissue for inclusion in the meta-analysis. The first selection criterion was sample size, i.e. if the number of analyzed samples was substantially (i.e. at least 30%) larger in one brain tissue versus the other, we retained the larger sample and excluded the other. Otherwise the prioritization on which brain tissue to include was based on the PD Braak staging (14) in order to maximize power (i.e. assuming that brain regions affected earlier in the disease course will show more pronounced effects). That is, the tissue from the region affected earliest in the disease process was selected for inclusion. To assess potential bias introduced by this “prioritization” strategy, we performed sensitivity analyses by including data from “lower priority” regions instead. For the other strata (blood-derived tissue and CSF), only one specimen subtype was assessed per study, thus prioritization was not applicable.

If a study reported several *p*-values for the same miRNA in the same samples based on different experimental or analytical methods (e.g. microarray versus RT-qPCR, different normalization approaches), we re-assessed whether one method was preferential to the other based on the information provided in the publication (e.g. higher accuracy/reliability), and only the most accurate result was included. If no decision could be reached, we chose a conservative approach and retained the largest *p*-value. For *p*-values reported with a reference to a predefined significance threshold only (applicable to data from 19/40 publications and a total of 121/2133 data entries), we used the following conservative conversions: “*p ≥* 0.05” and “p *≥* 0.01” were converted to “*p*=0.5”, *“p* <0.05” to “*p*=0.025”, “*p* <0.01” to “*p*=0.005”, “*p* <0.001” to “*p*=0.0005”, “*p* <0.0001” to “*p*=0.00005”. In three instances, the *p*-value in an article appeared as “0.0000”; this was converted to “0.00005”.

### Statistical analysis

#### Meta-analyses

Whenever possible, we calculated fixed-effect and random-effects (DerSimonian and Laird (15)) meta-analyses based on Hedges’ g as a standardized mean difference between idiopathic PD and unaffected control individuals. Depending on data reporting in the individual studies, Hedges’ g was calculated using different approaches: It was either calculated based on means and standard deviations or based on mean differences and corresponding measures of variance. In cases where no direct effect sizes were provided but the test statistic was described in sufficient detail, Hedges’ g was approximated from the reported test statistics as described previously (16). Whenever at least three independent datasets with Hedges’ g estimates were available effect-size based meta-analyses were calculated using the R package ‘meta’ (https://cran.r-proiect.org/web/packages/meta/meta.pdf). Between-study heterogeneity of effect-size based meta-analyses was quantified using the *I*^*2*^ metric, which represents the estimate of percentage of heterogeneity that is beyond chance.

In case data of effect-size estimates were not available we performed meta-analyses on provided *p*-values and directions of effects if these were available in ≥3 of independent datasets using a customized R script transforming *p*-values into signed z-scores using Stouffer’s method (17) (https://www.r-proiect.org; available upon request), similar to an approach described previously for the meta-analysis of genetic association data (18). This method allows to combine results even when effect size estimates and/or standard errors from individual studies are not available or are provided in different units (18). Briefly, the direction of effect and the *p*-value observed in each dataset were converted into a signed Z-score. Z-scores for each miRNA were then combined by calculating a weighted sum, with weights being proportional to the square root of the effective sample size for each dataset. The primary meta-analysis for each miRNA was calculated based on the fixed-effect model, and, if additional independent data were available, the *p*-value based model. Random-effects models were calculated for comparison to the fixed-effect model. For diverging results between the fixed-effect and the random-effects models, forest plots, heterogeneity estimates, and effect directions across datasets were further investigated. Significance was defined using Bonferroni correction for multiple testing. This was based on the number of the primary meta-analyses performed across all three specimen strata (i.e., *α*=0.05/160=3.13×10^-4^).

#### Classification of study-wide significant results according to the presence of heterogeneity

Study-wide significant meta-analysis results that showed no or little effect-size heterogeneity (in the effect-based meta-analyses) and/or consistent direction of effects across datasets (in the *p*-value based meta-analyes) were classified as showing “strong” support for a genuine involvement in PD. In the presence of heterogeneity, the random-effects model is considered more conservative than both the *p*-value based and fixed-effect methods. However, as the random-effects model also tends to “penalize” consistent results that show heterogeneity on the same side of the null, we re-investigated the forest plots of effect-size based meta-analyses in such cases. If study-wide significant meta-analyses showed effect-size heterogeneity due to variance of effect size estimates primarily at the same side of the null, we classified the respective miRNA as showing “strong” evidence and as “suggestive” if effect size estimates were on both sides of the null.

In cases where only *p*-value based meta-analyses could be performed due to a lack of sufficient effect-size data reported in individual studies, we also classified the consistency of effect direction using the above categories. Accordingly, miRNAs with meta-analysis results showing substantially differing directions of effect across independent datasets were labeled as providing “suggestive” and otherwise as providing “strong” evidence for an involvement in PD.

#### MiRNA target gene analysis

In order to assess indirectly whether any of the significantly differentially expressed miRNAs in brain may be involved in PD pathogenesis, we tested for a potential enrichment of their target genes in results of the latest genome-wide association study (GWAS) in PD (2,3). To this end, summary statistics from 7,773,234 single-nucleotide polymorphisms (SNPs) were obtained from PDGene (http://www.pdgene.org) (3), and analyzed using two different approaches for miRNA target site definition. Firstly, we downloaded human miRNAs and corresponding experimentally validated miRNA targets from MiRTarBase (v. 6.1; http://mirtarbase.mbc.nctu.edu.tw/) (19). We used MiRTarBase since it lists miRNA-target interactions reported in the literature that have been experimentally validated e.g. by reporter assay, western blot, microarray and/or next-generation sequencing experiments. Secondly, we used brain-specific miRNA-target gene interactions predicted with AGO2 HITS-CLIP miRNA data published by Boudreau et al. (20). To this end, we mapped Ensembl gene identifiers from the data of Boudreau et al (20) to EntrezGene identifiers based on Ensembl v. 87 (http://www.ensembl.org). The corresponding gene sets from MiRTarBase and Bouddreau et al. (20) were analyzed with Pascal (21) using 1000 Genomes samples (CEU) for assessment of linkage disequilibrium. Pascal combines SNP-based GWAS summary statistics to gene set scores and tests for enrichment of significant findings using a χ2 test and an empirical method.

In addition, we evaluated which top brain miRNAs bind to mRNAs from genes located in the established PD risk loci (2-4) (PD genes assigned for each locus according to Chang et al. (4)) and to the established causal PD genes *LRRK2, SNCA, VPS35, PRKN, PINK1*, and *PARK7* (a.k.a. *DJ1*) (6).

Furthermore, we evaluated whether any individual SNP (apart from the established, i.e. genome-wide significant, risk SNPs) located in the miRNA target genes (± 10 kb) was significantly associated with PD in the GWAS data (2,3). Adjustment for multiple testing was performed using Bonferroni correction for the number of tested target genes for all top miRNAs (i.e., *α*=0.05/532= 9.40×10^-5^). Finally, we investigated whether the respective PD-associated SNPs (or their proxies using a pairwise linkage disequilibrium estimate of r^2^>0.6 as threshold) may directly alter binding of the respective target miRNA(s). To this end, we mapped the SNPs and their proxies to the target sites of the top brain miRNAs as predicted by Targetscan (release 7.2; http://www.targetscan.org/vert_72/).

## RESULTS

### Description of eligible studies

The PubMed search yielded 599 publications, which were screened for eligibility of inclusion. A total of 52 publications were eligible for initial data extraction. After QC, data from 47 independent datasets across 40 publications were subsequently included in the meta-analyses. Reasons for the exclusion of eligible datasets from meta-analysis are summarized in Figure 1 and Table 1.

MiRNA expression data included in the meta-analyses were derived from brain tissue, CSF, and/or blood-derived samples. Eleven of the total of 47 datasets included in the meta-analysis were based on brain, 32 datasets on blood-derived samples, and four datasets on CSF. Only one of the included publications tested more than one of the three specimen types (blood and CSF (22)). Sampled brain regions of datasets included in the meta-analyses comprised substantia nigra/midbrain (n datasets=6), neocortex (n=4, comprising frontal, prefrontal, temporal, and anterior cingulate cortex), and amygdala (n=1; Table 1). The median number of study participants per dataset was 46 across all studies (interquartile range [IQR] 12-95, range 4-250) irrespective of the specimen type analyzed. The median number of individuals was 11 (IQR 8-16, range 4-62) for brain tissue, 81 (IQR 41-114, range 13-250) for blood-derived specimens, and 93.5 (IQR 70-115, range 58-122) for analyses of CSF. Twenty-seven out of the 47 independent datasets provided (albeit sometimes only for a subset of miRNAs) data that allowed for the calculation of Hedges’ g as the standardized mean difference.

Across all 40 studies included in the analyses presented here, half of the eligible studies (20/40, 50%) stated explicitly that they had performed age matching in their study design. Furthermore, information on the age distribution in patients and controls was provided for 26 datasets, and this distribution was comparable in most instances (average difference in patients and controls across all 26 datasets: 2.5 years, Supplementary Table 1). Four studies indicated statistically significant differences in the age distribution between patients and controls. Similar observations were made for the reporting of sex matching (40% report sex matching, average difference: 3.1%; Supplementary Table 1).

Twenty-two of all 40 studies used a targeted (“candidate miRNA”) approach to quantify miRNAs using RT-qPCR (n=20 studies), northern blotting (n=1), or a combination of methods (n=1). The remaining 18 studies applied a hypothesis-free (“mirRnome-wide”) screening approach using microarrays (n=5), next-generation sequencing (n=6), or TaqMan array micro RNA cards (n=7). The five studies using microarrays as an initial hypothesis-free approach applied targeted quantification methods for the top miRNAs in the same samples for validation.

The median number of miRNAs analyzed per study and included in the meta-analyses presented here was 3 (IQR 1-5) ranging from 1 to 123. Only four studies presented data on more than 100 miRNAs (Table 1). Overall, data for a total of 1,004 different miRNAs were reported across all studies, of which 140 had been assessed in at least three independent datasets in at least one specimen stratum and were thus eligible for meta-analysis (Supplementary Tables 2 and 3). Another 327 miRNAs had been assessed in two studies in at least one specimen type, and the remaining 537 had been assessed in only a single study in a single specimen type. Seventeen of the 140 miRNAs were meta-analyzed in both brain and blood strata, one miRNA was meta-analyzed in brain and CSF, and one miRNA in all three strata, overall resulting in 160 individual meta-analyses (Supplementary Tables 2 and 3, Figures 2 and 3).

**Figure 2.**
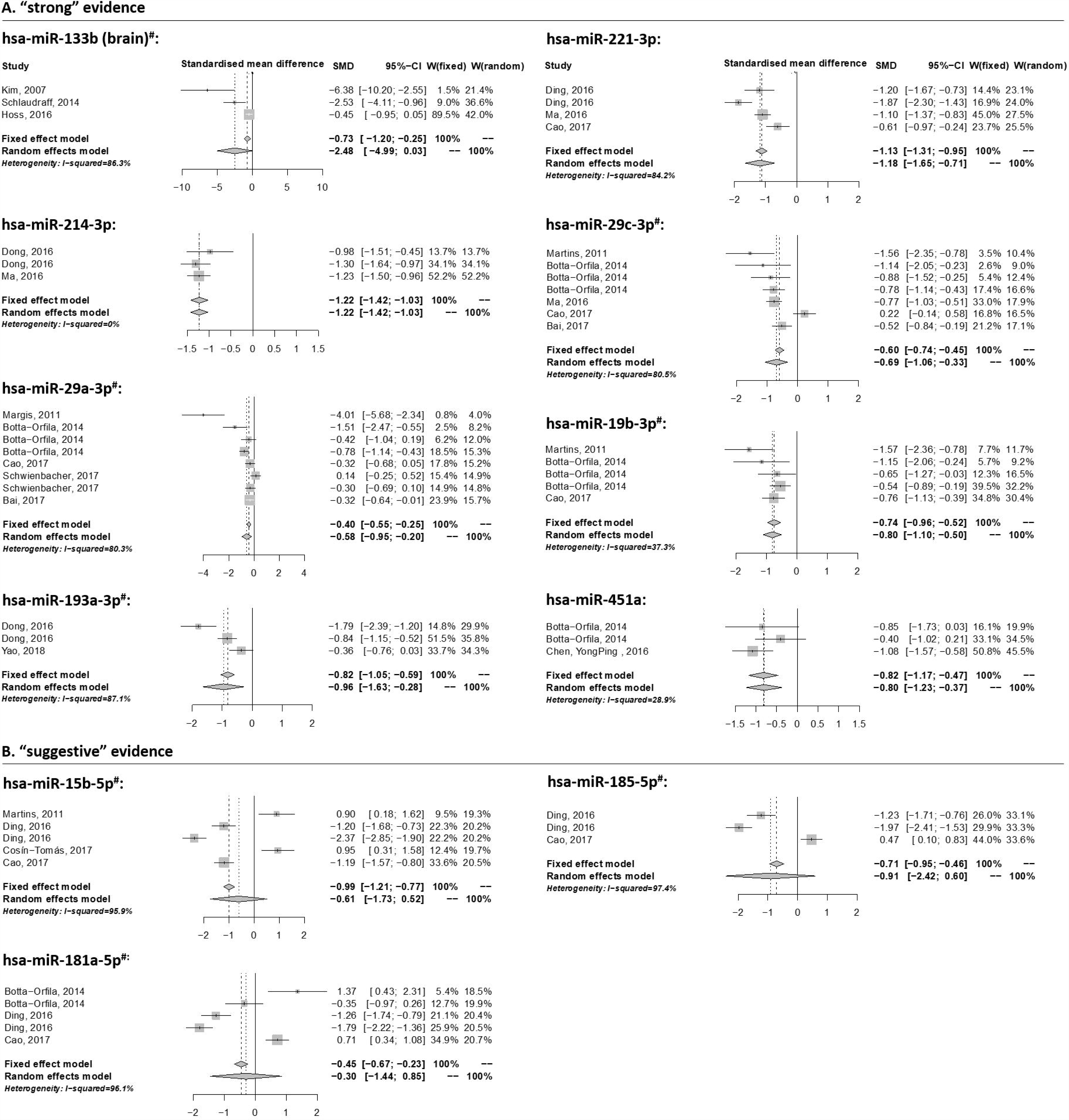
Forest plots of study-wide significant fixed-effect and random-effect meta-analyses on published miRNA expression data in idiopathic Parkinson’s disease and unaffected control individuals. Study-wide significant meta-analysis results (α=3.13×10^-4^) were classified as showing “strong” and “suggestive” evidence for differential expression in Parkinson’s disease according to heterogeneity assessments (see methods). Note that for several miRNAs, extended datasets were available for *p*-value based meta-analyses, which are therefore considered as primary meta-analyses (respective miRNAs are labeled with the symbol ^#^ also see Table 2 for details).

**Figure 3.**
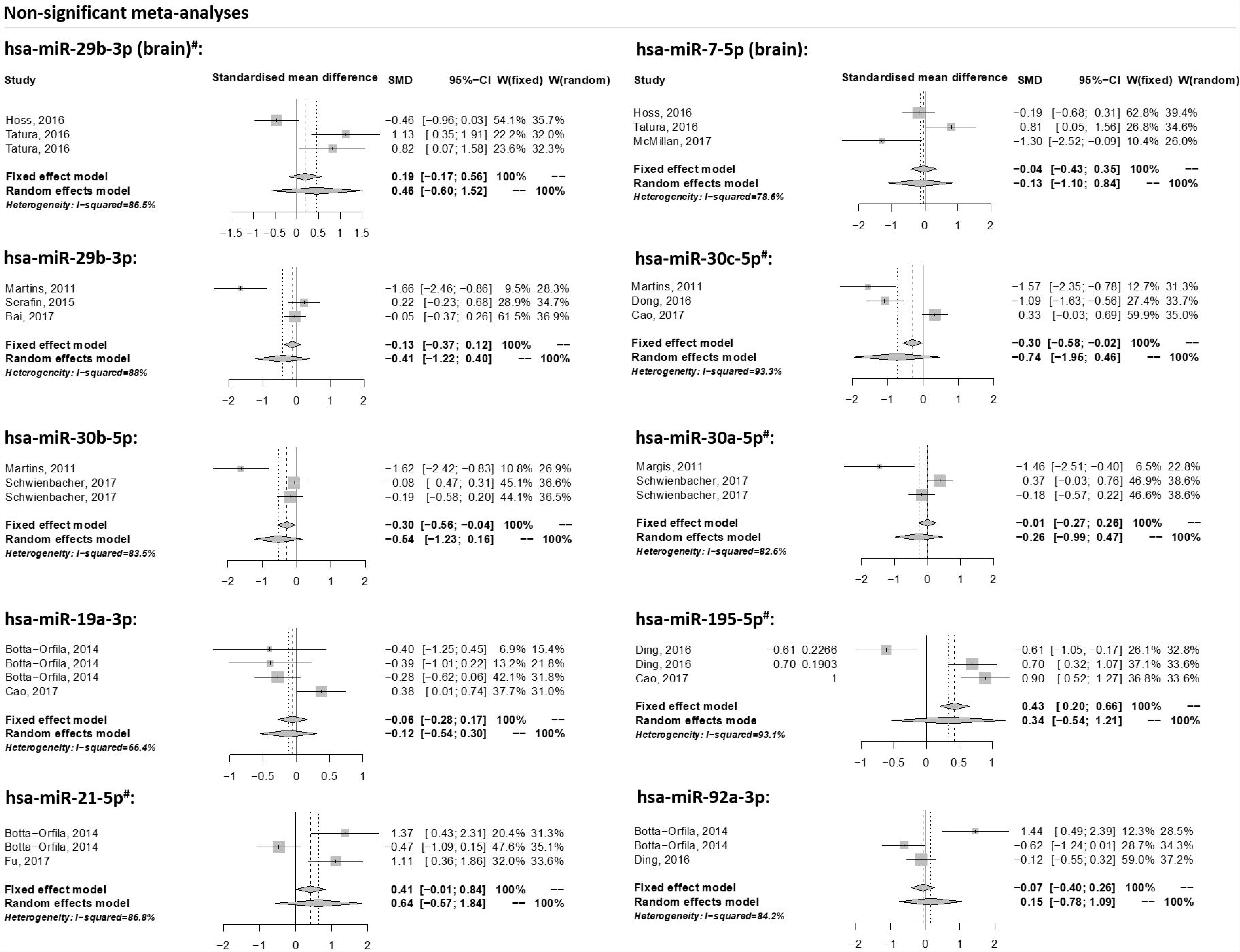
Forest plots of non-significant fixed-effect and random-effect meta-analyses on published miRNA expression data in idiopathic Parkinson’s disease and unaffected control individuals.

**Table 2.**
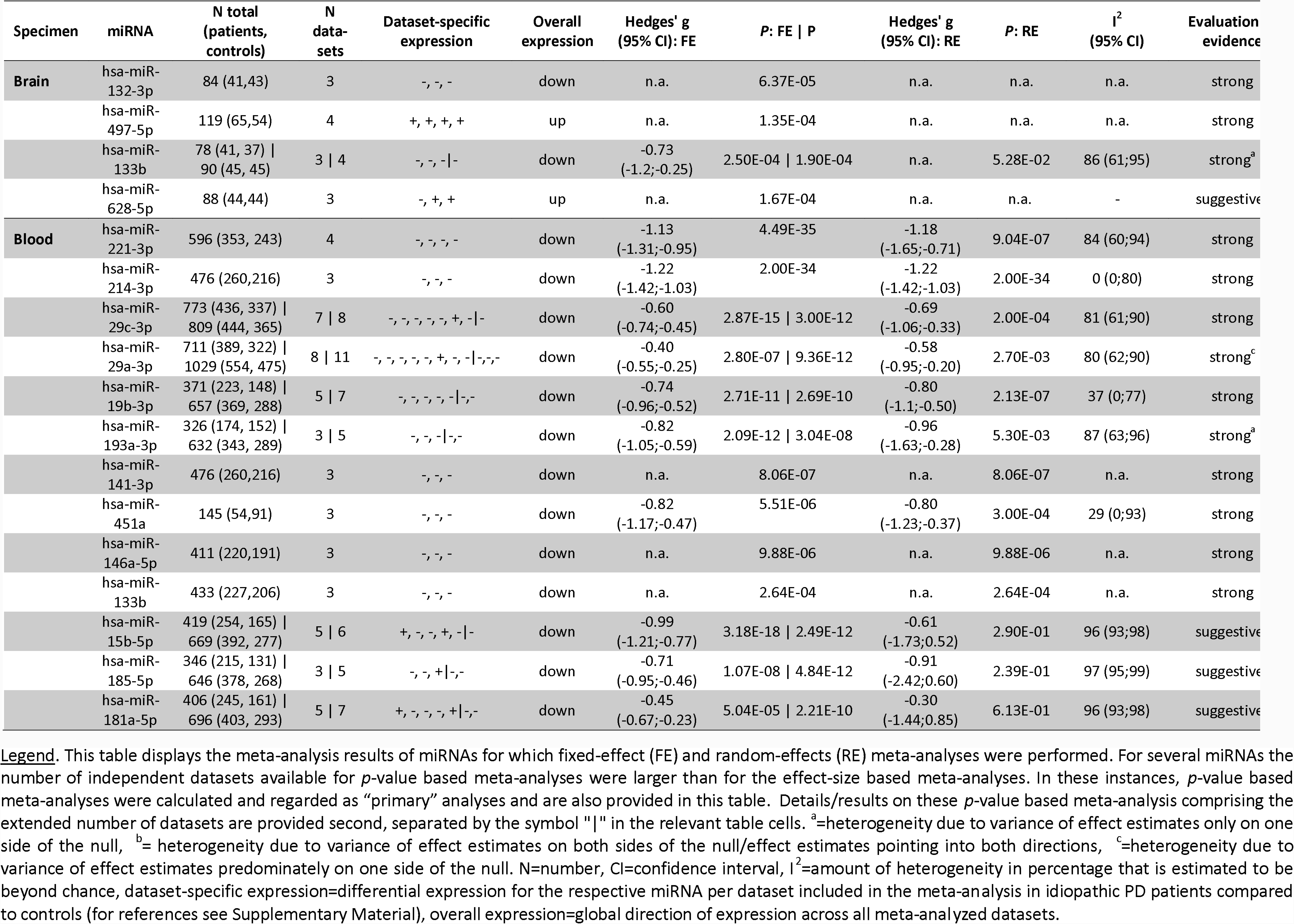
Significant meta-analysis results of differentially expressed miRNAs in brain and blood specimen of Parkinson’s disease patients and controls. *Note to the editor and reviewers: This table has been extensively modified, individual changes have not* been *highlighted*

### Meta-analysis results

One hundred twenty five meta-analyses were based on data collected in brain tissue, 31 in blood-derived samples, and four in CSF. Twenty-one of these meta-analyses were calculated based on effect sizes (Hedges’ g) using fixed-effect and random-effects models (brain: n=3, blood: n=18; Supplementary Table 2, Figures 2 and 3). For more than half of these meta-analyses (13 out of 21), additional datasets were available allowing extended meta-analyses based on *p*-values (Supplementary Table 2).

The median number of datasets included per primary meta-analysis across all miRNAs in brain, blood, and CSF was 3 (max. 4), 3 (max. 11), and 3 (max. 4), respectively. The median combined sample size across all miRNAs in brain, blood, and CSF was 88 (IQR 87-98), 339 (IQR 267-596), and 309 (IQR 309-323.5), respectively. On average, approximately equal numbers of patients and controls were included in each meta-analysis (Supplementary Tables 2 and 3).

Three of the 125 miRNAs meta-analyzed in brain showed study-wide significant (*α*=3.13×10^-4^) differential expression in idiopathic PD versus controls subjects with effect estimates pointing into the same direction of effect for each meta-analysis (classified as “strong” evidence, Table 2). One miRNA was up-regulated (hsa-miR-497-5p, *p*=1.35×10^-4^), while two (hsa-miR-132-3p, *p*=6.37×10^-5^, hsa-miR-133b, *p*=1.90×10^-4^) were downregulated in idiopathic PD compared to control subjects (Table 2). Furthermore, the meta-analysis result for one brain miRNA (hsa-miR-628-5p) reached study-wide significance in the *p*-value based model (*p*=1.67×10^-4^; effect-size estimates were not available), but effect directions were heterogeneous. Consequently, we classified this miRNA as showing “suggestive” evidence for differential expression (Table 2). In addition, 34 brain miRNAs showed nominally significant (*α*=0.05) differential expression (Supplementary Tables 2 and 3); however, these results did not survive multiple testing correction (*α*=3.13×10^-4^). Sensitivity analyses on the prioritization of multiple brain areas analyzed in the same samples showed that meta-analysis results were sufficiently robust regarding our prioritization procedure (Supplementary Table 4).

Ten out of 31 meta-analyzed miRNAs from blood-derived samples showed study-wide significant (*α*=3.13×10^-4^) differential expression in idiopathic PD versus control subjects (*p*-values ranging from 4.49×10^-35^ to 2.64×10^-4^) with effect estimates nearly always pointing into the same direction in each meta-analysis (“strong” evidence, Table 2). All ten miRNAs were down-regulated in idiopathic PD compared to control subjects (Table 2). The miRNA with the most statistically significant differential expression in blood was hsa-miR-221-3p (*p*=4.49×10^-35^). In addition, three miRNAs (i.e., hsa-miR-15b-5p, hsa-miR-185-5p, and hsa-miR-181a-5p) showed study-wide significant differential expression in blood specimen in the fixed-effect meta-analyses in the presence of substantial in-between study heterogeneity, i.e., with effect estimates on both sides of the null (Table 2). We therefore classified the results for these three miRNAs as “suggestive”. Seven additional miRNAs showed nominally significant (α=0.05) differential expression in the primary meta-analyses (Supplementary Tables 2 and 3), but did not survive multiple testing (α=3.13×10^-4^).

Of the four miRNAs meta-analyzed in CSF, none yielded significant results for differential expression in idiopathic PD versus control individuals (Supplementary Table 3).

Interestingly, hsa-miR-133b was study-wide significant in both brain (*p*= 1.90×10^−4^) and blood (*p*= 2.64×10^−4^) and was down-regulated in both specimen groups. Furthermore, miRNAs hsa-miR-19b-3p, hsa-miR-185-5p, and hsa-miR-29a-3p showed at least nominally significant expression differences in both brain and blood. Hsa-miR-19b-3p and hsa-miR-185-5p were down-regulated in both brain (*p*= 7.29×10^-4^and *p*=0.0034, respectively) and blood (*p*= 2.68×10^-10^ and *p*=4.84×10^-12^, respectively) in PD versus controls. Hsa-miR-29a-3p was up-regulated in brain (*p*=0.0322) and down-regulated in blood (*p*=9.36×10^-12^; Supplementary Tables 2 and 3).

### Target gene analysis of top differentially expressed brain miRNAs

We assessed whether SNPs in or near genes that represent targets of the top candidate brain miRNAs (three classified as showing “strong” and one as showing “suggestive” evidence) may also contribute to PD risk (regardless of whether they alter protein function or gene expression by non-miRNA mediated or by miRNA-mediated mechanisms). This may represent an independent line of evidence supporting a potential role of the respective brain miRNA(s) in PD pathophysiology. Based on published functional data available in miRTarBase (19) and on brain-specific HITS-CLIP data (20), three of the four brain miRNAs (showing “strong” [n=3] and “suggestive” [n=1] evidence for differential expression) were found to target mRNAs from genes located in established PD risk loci or from causal PD genes. For instance, based on the available brain HITS-CLIP data, hsa-miR-132-3p binds to the mRNAs of *SNCA* and of *SCN3A*, and hsa-miR-497-5p binds to the mRNA of *CCNT2* (Supplementary Table 5). *Note to the editor/reviewers: Supplementary Figure 1 removed as per journal policy which does not permit figures in the supplement.*

Considering all sets of genes targeted by any of the top four brain miRNAs, no set of targets showed significant enrichment (*α*=0.05) for genetic association with PD from GWAS data (Supplementary Table 6). However, the GWAS results of genetic variants mapping in target genes of the four brain miRNAs (after exclusion of the established risk loci already evaluated above) revealed nine additional loci that showed significant association with PD (*α*=9.40×10^-5^, Bonferroni-adjusted for the number of evaluated target genes [n=532], Supplementary Table 7).

Based on TargetScan predictions, a proxy (rs2977461) of one PD GWAS SNP (rs2944758, r^2^=0.62) is located 19bp downstream of the seed site of miR-132-3p (chr8:141541307-141541314) in the 3’UTR of *AG02* and may thus possibly affect the binding of this miRNA to its target.

### Comparison of miRNAs featured in original publications versus meta-analysis results

Across all eligible studies a total of 73 different miRNAs were “featured” in the original publications, i.e. they were prominently highlighted as showing differential expression in PD patients versus controls in the abstract of the respective publication. Only 8 (~11%; hsa-miR-1-3p, hsa-miR-7-5p, hsa-miR-30b-5p, hsa-miR-34b-3p, hsa-miR-146a-5p, hsa-miR-195-5p, hsa-miR-205-5p, hsa-miR-214-3p) of these were featured in two studies, and 6 (~8%; hsa-miR-19b-3p, hsa-miR-24-3p, hsa-miR-29a-3p, hsa-miR-29c-3p, hsa-miR-133b, hsa-miR-221-3p) in more than two studies. More than half of these featured miRNAs (45/73, 62%) were meta-analyzed in our study. Of note, 13 of these 45 miRNAs (~29%), indeed, showed study-wide significant association (α=3.13×10^4^, with “strong” and “suggestive” evidence) in our meta-analyses, whilst an additional ten (~22%) showed nominally significant association (*α*=0.05). In contrast, nearly half (i.e. 22 of 45 miRNAs (49%)) that had been prominently highlighted in at least one publication did not show any significant results in our meta-analyses. In addition, and perhaps more importantly, miRNAs miR-497-5p and miR-628-5p, showing “strong” and “suggestive” evidence for association, respectively, in our brain-stratified meta-analyses, and hsa-miR-451a, showing study-wide significance with “strong” evidence in the blood-stratified meta-analyses, were not featured in any of the original studies.

### Comparison of original versus replication evidence

To further assess the reproducibility of significant miRNA expression results, we compared all at least nominally significant *p*-values from the original study with results from independent replication data only (replication data were combined by meta-analysis, where applicable; Figure 4). For 34 (21%) of all 160 meta-analyses, nominally significant (two-sided *α*=0.05) differential miRNA expression was recorded by us for the first study. Less than half of these results (n=12, 35%) were replicated with at least nominal significance (one-sided *α*=0.05) when all available independent replication data were combined, and nine of these 12 results that replicated also yielded study-wide significance (two-sided *α*=3.13×10^-4^) upon meta-analysis of *all* data (i.e., combining original and replication data). Interestingly, the failure of replication of original results was predominately observed in CSF and brain while most blood-based findings showed good evidence for replication (Figure 4).

**Figure 4.**
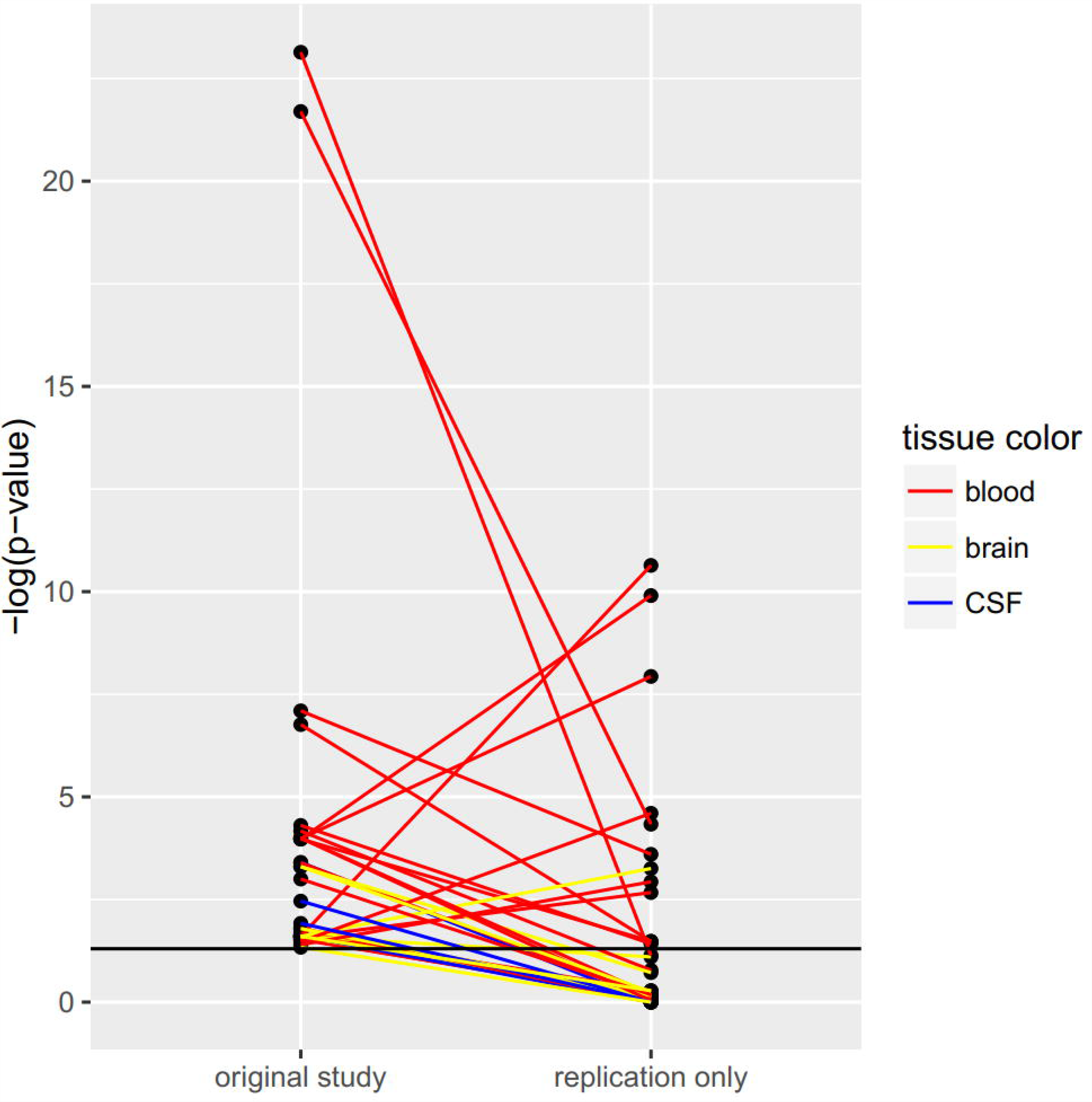
Comparisons of original and replication *p*-values. This figure displays all at least nominally significant two-sided *p*-values of the respective original studies (data from independent datasets derived from the original study were combined by meta-analysis where applicable), and the corresponding (one-sided) *p*-value from all replication data only (combined by meta-analysis where applicable). Note that *p*-values from all other meta-analyses in this paper are two-sided; a one-sided *p*-value was chosen here to take into account the directions of effect in the replication data. Corresponding *p*-values of original and replication data are connected by a line (yellow line = brain-stratified results, red line = blood-stratified results, blue line = cerebrospinal fluid-stratified results). The y axis shows the negative log of the *p*-value, i.e. larger values indicate more significant results. The horizontal black line corresponds to a *p*-value of 0.05.

## DISCUSSION

Following a systematic literature search and data extraction, we analyzed data from all hitherto published eligible miRNA expression studies in PD patients versus controls. We identified 17 miRNAs that were significantly differentially expressed in brain or blood across at least three independent studies. Based on heterogeneity assessments, we classified 13 of these miRNA as showing “strong” evidence for differential expression and four miRNAs as showing “suggestive” evidence. Interestingly, some of the top brain miRNAs target mRNAs of genes that are central in PD pathophysiology. The most compelling finding relates to miRNA hsa-miR-132-3p binding to the mRNA of *SNCA.* To the best of our knowledge, our study represents the first quantitative assessment of published miRNA expression data in PD. Furthermore, we are not aware of any other neurodegenerative research field having applied a comparable approach to collate published miRNA expression results of individual miRNAs by meta-analysis. We are aware of one publication (23) that meta-analyzed sensitivity and specificity estimates of miRNA profiles in Alzheimer’s disease in seven out of >100 differential miRNA epxression studies in that field. This approach, however, is substantially different from our approach, wich aims to pinpoint individual miRNAs differentially expressed between affected and unaffected individuals. Therefore, our study not only provides unique insights into the current knowledge of individual miRNA expression differences in PD but may also be taken as an example for performing equivalent analyses in other neurodegenerative diseases. In fact, our group is currently performing a similar field-wide analysis for differential miRNA expression in Alzheimer’s disease (AD). Preliminary results from that ongoing effort suggest that possibly up to three miRNAs (i.e. hsa-miR-29c-3p, hsa-miR-146a-5p, and hsa-miR-451a) showing differential expression in blood in PD, also appear to be differentially expressed in blood from AD patients when compared to controls (Takousis et al, in preparation).

One of the strengths of this study is the increase in sample size (and thus power) by combining all eligible data into one statistical test. As outlined above, sample sizes of individual miRNA studies are often small, especially in studies of brain tissue. By meta-analysis, we were able to increase the sample size substantially. In addition, errors occurring only in a single dataset will have a less pronounced impact on the resulting test statistic. Still, most of our brain-stratified meta-analyses (median n=88) are underpowered to detect only modest changes in miRNA expression. At the same time, significant results need to be considered with caution. Thus, a substantial increase in sample size should be one of the major objectives in future miRNA expression studies focusing on brain tissue.

Our study shows that the majority of miRNAs featured in the original publications or showing significant results in the first study cannot be replicated in independent investigations and do not have statistical support for differential expression in our meta-analyses. Along these lines, qualitative reviews on the role of miRNAs in PD are largely based on a (subjective) selection of the literature that does not hold up to systematic meta-analyses. For instance, in five recent articles reviewing the role of miRNAs in PD based on human expression or on experimental data (7,24-27) (including one systematic review (27)), 190 miRNAs were highlighted as being potentially relevant in PD (Supplementary 8). Of these, expression data were lacking or sparse for 113 (59%), i.e. they could not be meta-analyzed here. Among the remaining 77 miRNAs highlighted by at least one review, only 13 (7% of the 190 miRNAs) showed evidence for differential expression in PD in our meta-analyses. Furthermore, three of our top miRNAs (hsa-miR-146a-5p, hsa-miR-497-5p and hsa-miR-628-5p) were not mentioned in any of the five reviews. These observations highlight the need for independent replication and validation of proposed miRNAs as well as for regular quantitative - rather than merely qualitative - assessments of the available evidence in the literature.

Most of our significant results were based on blood expression data. While these results will likely not reveal novel insights into PD’s pathophysiology, these miRNAs may still have the potential to serve as “classification markers” for (prevalent) PD. It should also be noted that gene expression is not only tissue-specific but also variable over time. Thus, differential expression of miRNAs does not allow to draw conclusions on cause-effect relationships in PD. This is true for both blood and brain and for any investigation examining (prevalent) PD patients. In this context it is noteworthy that all eleven miRNAs in the blood-based results appear to be “downregulated” in idiopathic PD as compared to control subjects. This may reflect changes in gene expression and/or cell compositions as a result of disease progression or maybe most likely treatment effects. Further, in the brain-derived results, especially those from substantia nigra, it is also possible that expression differences might only reflect changes of cellular composition in the diseased tissue. As most studies normalize the results using general house-keeping genes, such effects will not necessarily be removed entirely. An alternative way to quantify miRNA expression would be to perform single-cell experiments in cells of interest, e.g. dopaminergic neurons. However, while a meta-analysis has recently been published for mRNA-based transcriptomics studies applying laser capturing for single cell analysis in the substantia nigra (28), equivalent data on miRNAs are currently too sparse.

Furthermore, most publications do not provide any information on disease duration, severity, and treatment of patients, and, for brain tissue, neuropathological progression markers. Thus, the impact of these factors on the respective miRNA results is impossible to assess adequately. In addition, a study design that does not consider age and/or sex matching for patients and controls may produce biased gene expression results. As described in the results section, the majority of datasets had comparable age and sex distributions in patients and controls. Notwithstanding we cannot exclude that missing age and/or sex matching has had an impact on some of our meta-analysis results. Furthermore, we note that other variables such as the use of different eligibility criteria and recruitment schemes and diverse specimen retrieval protocols, as well as different methods of RNA extraction, miRNA expression measurements, and different statistical methods, etc, may impact the results of any individual study and may, thus, be one of the causes of in-between study heterogeneity. However, the current number of independent individual datasets per miRNA is too small to investigate the impact of these variables systematically, e.g. by performing sensitivity or meta regression analyses.

In the context of controlling the potential impact of external variables and in order to disentangle cause-effect relationships, the conduct of differential expression studies in animal models may be useful despite the potential limitations in translating these findings to the human system. To this end, we investigated (by screening titles and abstracts) how many of the 547 papers excluded from our systematic review would qualify for a similar meta-analytical approach based on animal models. This resulted in 11 differential miRNA expression studies in PD animal models. However, these studies were very heterogenous in that they investigated a range of different animals (mouse=6 studies, rat=2, drosophila=2, C.elegans=1) and focused mostly on non-overlapping miRNAs. Therefore, while these results could be regarded as informative in the context of our study, the currently available published data is too sparse to yield meaningful meta-analyses or robust qualitative assessments.

In this study, whenever possible, we applied effect-size based meta-analyses (n=21) based on the standardized mean difference (Hedges’ g), which allowed us to also quantify effect-size heterogeneity across the included datasets. Importantly, this list comprised 11 of the 17 miRNAs showing Bonferroni-corrected significance (with “strong” and “suggestive” evidence) in our primary meta-analyses. However, about 40% of all eligible studies did not provide precise effect estimates and/or variances and, thus, precluded the calculation of Hedges’ g. In cases where effect-size based meta-analyses were not possible (n=135) or where data were available for additional datasets on the same miRNAs (n=13/21 effect-size based meta-analyses), we performed systematic *p*-value based meta-analyses to collate the available published data. This is an established method often applied in the GWAS field (18) and the *p*-values of our fixed-effect meta-analyses corresponded well to those of the *p*-value based meta-analyses on identical sets of data (Supplementary Table 9). However, using *p*-values only does not allow to estimate the magnitude of gene expression differences, to quantify the heterogeneity of estimated effect sizes, or to perform additional analyses such as testing for small-study effects, which can be indicative of publication or selective reporting bias (29). However, except for the heterogeneity assessments none of these additional analyses were possible for our effect-size based data due to a lack of sufficient data. As proxy for in-between study heterogeneity for the *p*-value based meta-analyses, we assessed the consistency of effect directions qualitatively across individual datasets for study-wide significant miRNAs. This revealed potential evidence for heterogeneity, i.e. effect directions pointing in both directions of the null, for hsa-miR-628-5p expression in brain, and for to hsa-miR-15b-5p, hsa-miR-185-5p and hsa-miR-181a-5p in blood, for which heterogeneity was already observed in the effect-size meta-analyses for a smaller subset of data. Due to this heterogeneity, we classified the overall evidence for differential expression for these four miRNAs as “suggestive” only (Table 2).

Importantly, a proportion of publications (applicable to data from 19/40 publications and a total of 121/2133 data entries) did not report full *p*-values but reported them as “less than” or “greater than” a certain significance level. Here, we chose a conservative approach for including such data in our analyses (see methods section). Furthermore, the quality of our analyses can, at best, only mirror the quality of the underlying publications from which data were extracted. We performed a range of quality control checks to detect inconsistencies within studies, but cannot exclude that all errors were detected by this procedure. However, we do not expect any *systematic* error arising from errors and mistakes that may have remained undetected in the original publications. Nevertheless, these observations clearly highlight the need for a standardized and more transparent reporting of applied methodology, statistics and results in miRNA expression studies (30).

One additional limitation in combining data from the published domain is the potential presence of publication bias and/or selective reporting bias. Due to the lack of consistently reported effect size estimates in a part of eligible publications (see above), we were not able to assess potential hints for this bias quantitatively (e.g. by regression analyses (31)). To address this concern, we evaluated each publication for evidence that only a subset of the generated expression results were reported in detail (Supplementary Table 10). For nearly two thirds of all publications (i.e., 25/40, 63%) we did not find evidence for selective reporting of expression results. Fifteen publications had generated more data than provided in the publication. Five of these studies provided the identifiers of the miRNAs for which detailed results were not provided. This list contained ten of the 13 miRNAs differentially expressed in blood (with strong or suggestive evidence) according to our meta-analyses. This was due to few large-scale studies that had only highlighted the most prominent miRNAs. Meta-analyses in other fields (e.g. cancer) of miRNA and other regulatory RNA associations have pointed out the surprisingly high proportion of reported statistically significant results, which may be an indication of excess significance due to selective reporting (32,33). This pattern was not as prominent in the studies that we analyzed, where 15 of the identified studies (Table 1) did not feature any particular miRNAs eventually. In summary, we cannot exclude that selective reporting has inflated some of our meta-analysis results. Especially the blood-based meta-analysis results need to be considered with caution and warrant independent replication.

In conclusion, by systematically combining data from all eligible miRNA expression studies published to date, we identified 13 miRNAs that were consistently differentially expressed in PD patients and controls in brain or blood. Future studies will need to increase the sample size for miRNA-based studies on brain tissue. Our study is the first to compile published miRNA expression data in the field of neurodegenerative diseases in a systematic and standardized way. Thus, it may serve as a model for combining these data in other related fields.

## ACKNOWLEDGMENTS

We thank Lukas Duchrow and Colin Schulz for excellent assistance in the bioinformatic analyses. We also thank Fabian Kilpert for help with the figures. C.M.L. and L.B. received funding from the German Research Foundation (DFG; FOR2488/1: GZ LI 2654/2-1 and BE 2287/5-1), the Possehl Foundation, the Renate Maaß Foundation, and the University of Luebeck (section of medicine, J21-2016). J.S. was supported by the MD thesis research scholarship “Exzellenzmedizin” of the University of Luebeck. I.W. was supported by the Peter and Traudl Engelhorn Foundation.

## AUTHOR CONTRIBUTIONS

Study design: J.S., P.T., R.P., L.B., C.M.L., literature search and data extraction: J.S., P.T., I.O.G.I., V.D., statistical analyses/advice: J.S., I.W., G.R., H.B., J.P.I., C.M.L, writing of the manuscript: J.S., P.T., L.B., C.M.L., critical revision of the manuscript: all co-authors

